# Mapping diversification metrics in macroecological studies: Prospects and challenges

**DOI:** 10.1101/261867

**Authors:** Julián A. Velasco, Jesús N. Pinto-Ledezma

## Abstract

The intersection of macroecology and macroevolution is one of the most active research areas today. Macroecological studies are increasingly using phylogenetic diversification metrics to explore the role of evolutionary processes in shaping present-day patterns of biodiversity. Evolutionary explanations of species richness gradients are key for our understanding of how diversity accumulated in a region. For instance, the present-day diversity in a region can be a result of *in situ* diversification, extinction, or colonization from other regions, or a combination of all of these processes. However, it is unknown whether these metrics capture well these diversification and dispersal processes across geography. Some metrics (e.g., mean root distance -MRD-; lineage diversification-rate -DR-; evolutionary distinctiveness -ED-) seem to provide very similar geographical patterns regardless of how they were calculated (e.g., using branch lengths or not). The lack of appropriate estimates of extinction and dispersal rates in phylogenetic trees can limit our conclusions about how species richness gradients emerged. With a review of the literature and complemented by an empirical comparison, we show that phylogenetic metrics by itself are not capturing well the speciation, extinction and dispersal processes across the geographical gradients. Furthermore, we show how new biogeographic methods can improve our inference of past events and therefore our conclusions about the evolutionary mechanisms driving regional species richness. Finally, we recommend that future studies include several approaches (e.g., spatial diversification modelling, parametric biogeographic methods) to disentangle the relative the role of speciation, extinction and dispersal in the generation and maintenance of species richness gradients.

## Introduction

The causes of spatial variation of biodiversity are one of the most fundamental questions in ecology, biogeography and macroecology (Brown 1995, 2014, Brown and Lomolino 1998, Hawkins et al. 2012, Fine 2015, Jablonski et al. 2017). Current studies are integrating in a single framework the ecological and evolutionary mechanisms driving regional species diversity (McGaughran 2015, Pärtel et al. 2016, Cabral et al. 2017, Leidinger and Cabral 2017). However, only three macroevolutionary processes ultimately can modify the number of species in a region: speciation, extinction and dispersal (Wiens 2011, Fine 2015, Jablonski et al. 2017) (Figure 1). These processes can be modulated by species’ traits varying within clades (Paper et al. 2016, 2017, Jezkova and Wiens 2017, Moen and Wiens 2017), age of region (e.g., time-for-speciation effect; Stephens & Wiens, 2003), geographical area (Losos and Schluter 2000), or climatic conditions (Condamine et al. 2013, Lewitus and Morlon 2017).

**Figure 1.**
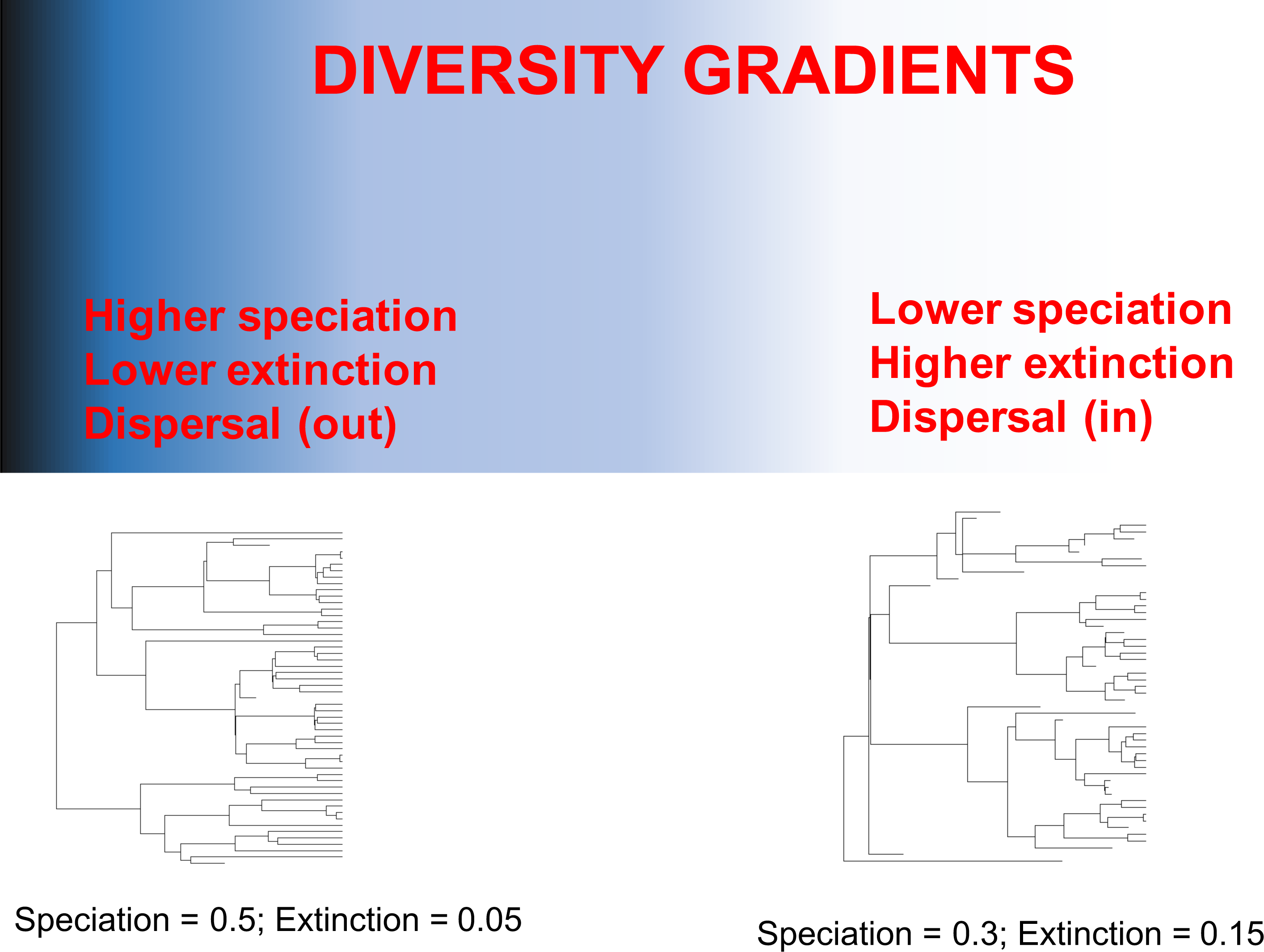
Diagram illustrating how differences in speciation, extinction, and dispersal rates between regions can generate a geographical species richness gradient. The phylogenetic trees below illustrate how the differences in speciation and extinction rates between two regional assemblages can shape a gradient of species richness (degraded blue colour).

The integration of different disciplines such as molecular phylogenetics, palaeontology, and historical biogeography have allow to infer a series of macroevolutionary processes across geography (Diniz-Filho et al. 2013, Fritz et al. 2013). It is well-known that fossil information is key to estimate with high confidence rates of speciation and extinction (Sepkoski 1998, Foote 2000, Quental and Marshall 2010, Rabosky 2010b). New methods correcting for sampling bias are able to generate improvements in the estimates of the speciation, extinction and net diversification rates (Silvestro et al. 2014, 2016). However, the causal mechanisms that underlying the geographical diversity gradients only can be established with greater confidence for a few taxonomic groups with adequate fossil record, such as marine bivalves (Jablonski et al. 2006, 2017), mammals (Silvestro et al. 2014) or plants (Antonelli et al. 2015).

As fossil data is not available or incomplete for most extant groups, model-based approaches used to estimate speciation and extinction rates in palaeontology were adapted to study the macroevolutionary dynamics using phylogenetic information (Nee et al. 1994, Morlon et al. 2010, Stadler 2013). Molecular phylogenies are becoming essential to the study of diversification dynamics across temporal and spatial scales for extant taxa (Wiens and Donoghue 2004, Rabosky and Lovette 2008, Stadler 2013, Morlon 2014, Schluter and Pennell 2017). Therefore, it is possible to reconstruct past diversification process based on the branching events of a phylogeny using a set of birth-death models (Nee et al. 1994, Nee 2006, Morlon et al. 2010, Stadler 2013, Morlon 2014). These birth-death models allow infer either a homogeneous process for an entire clade (Nee et al. 1994, Magallón and Sanderson 2001) or a heterogeneous process varying in time or in specific subclades of a tree (Paper et al. 2006, Rabosky and Lovette 2008, Alfaro et al. 2009). However, these birth-death models only account for temporal variation of the macroevolutionary processes and how translate these processes to the geography is still a matter of debate.

Macroecological studies use two main approaches to link the estimates of diversification with the geographical ranges of species (Hawkins et al. 2007, Algar et al. 2009, Qian et al. 2014, Pinto-Ledezma et al. 2017, Velasco et al. 2018). The first one uses a set of phylogenetic metrics as a proxy to capture the geographical signature of lineage diversification dynamics (Diniz-Filho *et al.*, 2013; Fritz *et al.*, 2013; Table 1; Figure 2). These metrics provide either an estimate of a per-species rate of diversification (e.g., mean root distance –MRD-, residual phylogenetic diversity –rPD-, mean diversification rate –MDR-), the phylogenetic structure of regional assemblages (e.g., phylogenetic species variability –PSV-) or the average age of co-occurring lineages in a given area (e.g., mean age; see Table 1). Each metric is calculated for each species in the phylogeny; therefore, we can associate the species’ values to its corresponding geographical range and generate a map with average values for cells or regions. Although these phylogenetic metrics only account for speciation events, macroecologists have used these maps as a proxy to test some evolutionary-based hypothesis in macroecological research (Diniz-Filho *et al.*, 2013; Fritz *et al.*, 2013; see Table 2 for a compendium of these hypotheses).

**Table 1:** Phylogenetic metrics and explicit diversification approaches used in macroecological studies to address evolutionary questions related with the geographical diversity gradients.

| Metric | Author | Description | Software / R package |
| --- | --- | --- | --- |
| MRD | Kerr and Currie 1999 | MRD is calculated by counting the number of nodes separating each terminal species in a regional assemblage or cell from the tips to root of the phylogenetic tree. This metric does not need that trees be ultrametric or have branch lengths. | metricTester |
| PD (residual) | Faith 1992 | PD is calculated by summing all the branch lengths of species co-occurring in a regional assemblage or cell. Residual PD is obtained from an ordinary least square regression between PD and species richness. | picante, metricTester, pez |
| PSV | Helmus et al. 2007, Algar et al. 2009 | PSV is calculated from a matrix where their diagonal elements provide the evolutionary divergence (based on the branch lengths) of each terminal species from the root to the tips of the tree, and the off-diagonal elements provide the degree of shared evolutionary history among species. Values close to zero indicates that all species in a regional assemblage or cell are very close related whereas values close to one indicate that species are not related. | picante, metricTester, pez |
| mean DR | Jetz et al. 2012 | DR is calculated as the inverse of a measure of evolutionary isolation (Redding & Mooers 2006) which sum all the edge lengths from a species to the root of the tree. The inverse of this evolutionary isolation metric therefore capture the level of splitting rate of each species (i.e., its path to a top). | FiSSE |
| Mean age | Latham and Ricklefs 1993 | The mean age of co-occurring species in a regional assemblage or cell simply is calculated tallying the age of each most recent common ancestor (MRCA) for each species and the averaged. | None |
| GeoSSE | Goldberg et al. 2011 | The geographic state speciation and extinction -GeoSSE- model is a trait-dependent diversification method linking geographic occurrence with diversification rates. These method allow to infer both speciation and extinction rates as movement (dispersal) rates among two regions. | Diversitree R package |
| BAMM | Rabosky 2014 | BAMM is a method that attempt to identify whether a phylogeny exhibit a single or various macroevolutionary regimes (i.e., different diversification dynamics). As speciation, extinction and net diversification rates are considered to be heterogeneous across the phylogeny it is possible to estimate a rate for each branch or species in the tree. | BAMM software and BAMMtools R package |

**Figure 2.**
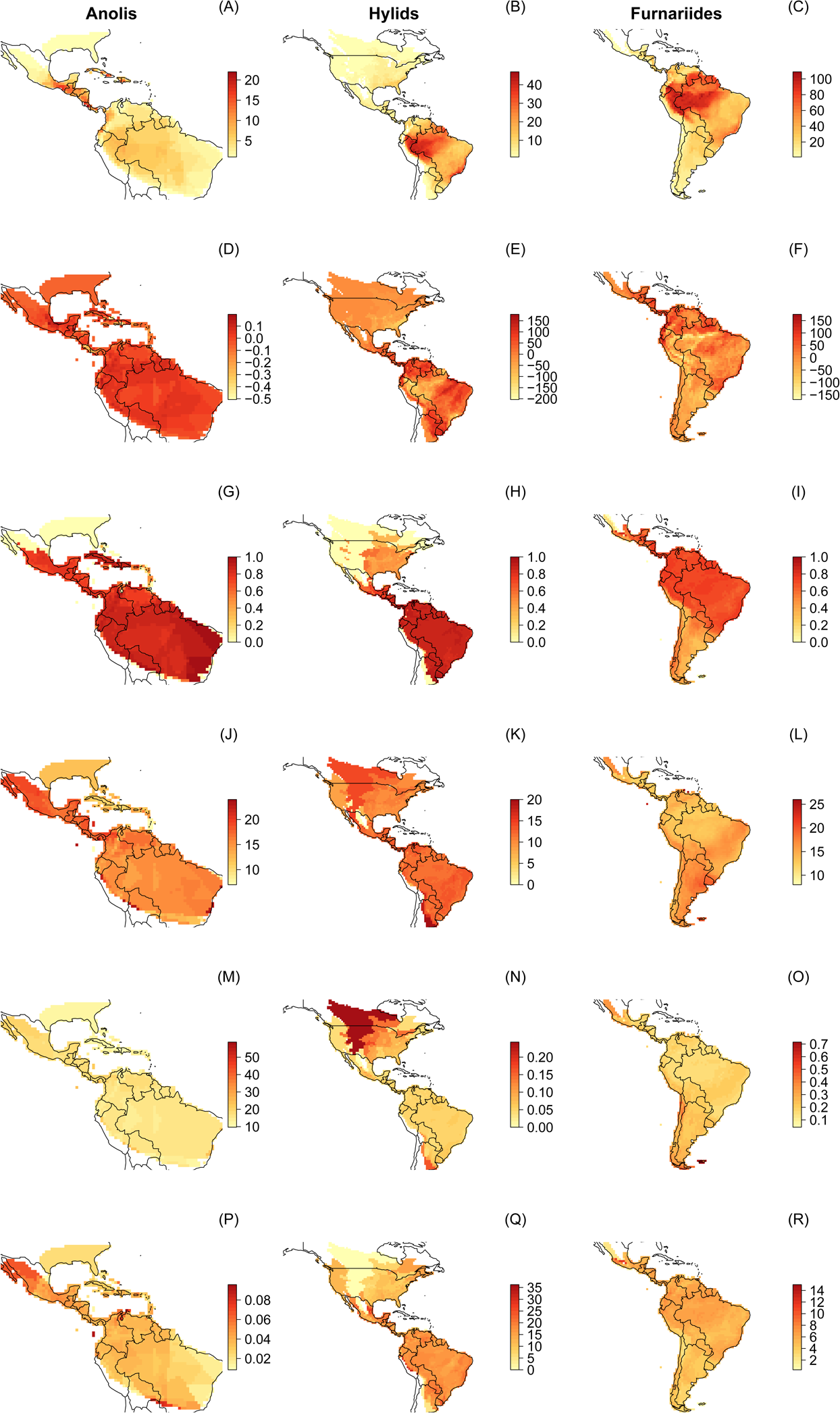
Geographical patterns of some phylogenetic metrics used in macroecological studies to explore evolutionary process underlying geographical diversity gradients (see also Table 1 for a detailed explanation). Left column *Anolis* lizards; Middle column: Hylid frogs; Right column: Furnariides birds. (A-C) observed richness patterns; (D-F) rPD: residual phylogenetic diversity (i.e., after controlling for species richness); (G-I) PSV: phylogenetic species variability; (J-L) MRD: mean root distance; (M-O) MDR: mean diversification rate; (P-R) Mean ages: average ages of species.

The second approach used by macroecologists consists in the explicit estimation of diversification parameters across geography (Goldberg et al. 2005, 2011, Ramiadantsoa et al. 2017). For instance, the geographic state speciation and extinction model –GeoSSE (Goldberg et al. 2011; Table 1) allows estimating speciation, extinction and dispersal parameters across two regions. It is possible to disentangle the relative role of each one of these processes on the generation and maintenance of the geographical diversity gradients (Rolland et al. 2014, Pulido-Santacruz and Weir 2016, Pinto-Ledezma et al. 2017). In addition, a recently developed Bayesian approach (BAMM; Rabosky 2014, Rabosky et al. 2014) allows to infer the balance of speciation and extinction in the generation of these biodiversity gradients (Rabosky et al. 2015, Sánchez-Ramírez et al. 2015, Morinière et al. 2016, Pinto-Ledezma et al. 2017). The BAMM approach allows both the inference of macroevolutionary dynamics for an entire clade (i.e., a macroevolutionary regime; Rabosky 2014) and also get estimates of per-species diversification rates (i.e., as a phylogenetic metric; Rabosky 2016) that can be mapped in a geographical domain. Although all these methods aim to obtain a geographical picture of the diversification processes, it remains unexplored if they can effectively capture these dynamics across regions.

In this paper, we conducted a review on macroecological literature to evaluate how evolutionary and biogeographic processes contribute to shape geographical species richness gradients. We review only those papers that make explicit use of phylogenetic metrics and/or explicit diversification approaches (Table 1). We divided our review in three main sections. In the first one, we discuss how studies use phylogenetic metric to test some evolutionary-based hypotheses underlying geographical diversity gradients and we explore some limitations of these metrics (see also Table 2). Also, we discuss to what extent these metrics are able to capture macroevolutionary dynamics in a spatial explicit context. We illustrate these using three case studies (Furnariides birds, Hylid frogs, and *Anolis* lizards; Figure 1) and explore how another approaches (e.g., diversification modelling and biogeographical approaches) can complement our inferences about diversification process across geography. In the second section, we discuss how dispersal and extinction processes are limiting these diversification inferences and we propose some research avenues to attempt to solve these problems. Using an explicit biogeographical approach, we test the role of dispersal on the geographical species richness patterns of the three case studies. Finally, in the third section, we call for the adoption of complementary approaches (e.g., extensive simulations, parametric biogeographical methods) in macroecological research with the aim to evaluate the relative role of speciation, extinction and dispersal process driving geographical biodiversity gradients.

**Table 2.** Description of some evolutionary hypothesis tested in macroecological studies as causal mechanisms of regional species richness.

| Hypothesis | References | Description | Predictions | Metrics and/or methods used to test | Limitations |
| --- | --- | --- | --- | --- | --- |
| Phylogenetic niche conservatism (PNC) | Wiens and Graham 2005 | Phylogenetic niche conservatism is the tendency of related species to inherit niche requirements from its the most recent common ancestors (Wiens & Graham 2005). | PNC predicts that regions where a clade originated will accumulate more species simply due to more occupation time and diversification rates tend to be similar between regions. The tropical niche conservatism hypothesis (TNC; Wiens and Donoghue 2004) is based on PNC to explain differences in species richness in tropical and temperate regions. | MRD, Mean age, GeoSSE, BAMM | 1) MRD metric fails to capture spatially dynamics of the balance of speciation and extinction and it is very hard to establish whether species richness in a region is only generated by higher speciation rates. Furthermore, MRD does not capture dispersal dynamic across regions and species richness in a given region can be generated from only dispersals from nearby regions (e.g., macroevolutionary sinks; CITA). 2) Mean age provide partial is able to test the role of PNC on geographical species richness because only it is possible to establish which regions have, in average, old clades and this not reflects whether many speciation events occurred there. 3) GeoSSE is potentially the only one approach that allow to disentangle these three process but it is only limited to two regions (e.g., tropical vs. temperate). In addition, GeoSSE has been criticized due its low statistical power (see Rabosky and Goldberg 2015). |
| Regional diversification (RD) | Buckley et al. 2010 | Differences in the balance of speciation and extinction across geography can explain differences in species richness between regions. | RD predicts that regions with striking differences in species richness are due to differences in macroevolutionary dynamics between regions. | residual PD, GeoSSE, BAMM | 1) Residual PD can be used to discriminate regions with rapid and slow diversification based on the expected phylogenetic diversity given species richness (Buckley et al. 2010). However, this metric ignores the contribution to dispersal to PD in a given region or cell. 2) GeoSSE can estimate speciation, extinction and dispersal rates between regions but again is limited to two regions. 3) BAMM potentially could be used to estimate speciation rates for regional clades but this method is unable to estimate dispersal rates between regions. |
| Out of the tropics (OTT) | Jablonski et al. 2006 | Species were generated in the tropical regions and dispersed to extratropical regions but maintain its presence in its ancestral areas | High rates of speciation are predicted in tropical regions in contrast with temperate regions. Asymmetric dispersal have occurred along the biogeographical history of a taxa from tropical to temperate areas. | MRD, Mean age, GeoSSE | These metrics are the same used to test the PNC/TNC hypothesis as we discuss above. |
| Time for speciation effect (TEE) | Stephens and Wiens 2003 | Tropical regions accumulated more species because their clades had more time to speciate than temperate regions. | Regions recently colonized had lower species richness than regions where clades colonized very early in the history of a clade. | Mean age | 1) Mean age does not provide an accurate description of which lineages colonized first a region. To test this hypothesis, it might be necessary to perform an ancestral range reconstruction of all co-occurring clades and estimate its diversification rates (i.e., total diversification for each independent colonized clade; Rabosky 2009; 2012). |

## Literature Review

We conducted a literature search in Web of Science for studies that explicitly have addressed questions on how speciation, extinction and dispersal have shaped geographical species richness gradients. We selected those papers that used either phylogenetic metrics (e.g., mean root distance -MRD-, phylogenetic diversity -PD-, phylogenetic species variability -PSV-; diversification rate -DR-; mean Ages; Table 1) or explicit macroevolutionary approaches (e.g., GeoSSE, BAMM; Goldberg *et al.*, 2011; Rabosky 2014). We compiled a list of 44 papers (Table A1), but we are aware that this likely is not an exhaustive search. The majority of papers reviewed are testing historical process shaping latitudinal diversity gradients (LDG) in various taxa.

## Testing Evolutionary Hypothesis Using Phylogenetic Metrics and Explicit Diversification Approaches

Here, we discuss how phylogenetic metrics are used to test evolutionary hypotheses related to the generation and maintenance of geographical diversity. Several studies used the mean root distance -MRD-metric to evaluate whether regional assemblages are composed of “basal” or “derivate” linages. First, this terminology should be avoided because it provides an incorrect interpretation of the phylogenetic trees (Baum et al. 2005, Crisp and Cook 2005, Omland et al. 2008). Although this metric does not incorporate information from branch lengths (Algar et al. 2009, Qian et al. 2015), it does provide the average number of nodes separating each species in a given region from the root of the phylogeny (Kerr and Currie 1999). MRD therefore provides information about the number of cladogenetic events (splits) that have occurred through the history of co-occurring lineages in each region (Pinto-Ledezma et al. 2017, Velasco et al. 2018). Under this view, MRD should be interpreted as a metric of total diversification (Rabosky 2009), where high MRD values indicating regional assemblages dominated by extensive cladogenesis and low MRD values indicating assemblages with few cladogenetic events. A main concern with this metric concern with the fact that it does not provide any information about what macroevolutionary dynamics have taken place in a region. For example, it is very hard to establish whether MRD allows to distinguish between diversity-dependent (Rabosky 2009, Rabosky and Hurlbert 2015) or time-dependent (Wiens 2011, Harmon and Harrison 2015) processes dominating regional diversity. Although the distinction between these two dynamics, and its relationship with the origin and maintenance of regional diversity, is an intense topic in the macroevolutionary literature (Rabosky 2009, 2013, Wiens 2011, Cornell 2013, Harmon and Harrison 2015, Rabosky and Hurlbert 2015). However, more empirical and theoretical work is necessary to establish what scenario plays a significant role in regional species richness assembly (Rabosky 2012, Etienne et al. 2012, Valente et al. 2015, Graham et al. 2018). It might worth to establish whether local ecological process scaling up to regional scales or emergent effects (i.e., the existence of a strong equilibrium process) governed the build-up of regional diversity (Cornell 2013, Harmon and Harrison 2015, Rabosky and Hurlbert 2015, Marshall and Quental 2016).

The time-for-species effect hypothesis state that the regional build-up of species richness is directly proportional to the colonization time of its constituent clades (Stephens and Wiens 2003b; Table 2). However, many phylogenetic metrics used to test this hypothesis, did not incorporate any age information (Fritz and Rahbek 2012, Qian et al. 2015). Qian et al. (2015) did some additional predictions for the time effect hypothesis regarding the phylogenetic structure of regional assemblages. We suggest that these predictions are not easily deduced from the original statement of the time-for-speciation effect hypothesis (Stephens & Wiens 2003). For instance, Qian et al. (2015, p. 7) predicted that regions with low species richness (e.g., extra-tropical regions) should be composed of more closely related species than regions with high species richness (e.g., tropical regions). This assumes that regions with low species richness were colonized recently and therefore these lineages had little time for speciation. However, it is also plausible consider that high extinction occurred in these poor species richness regions by marginal climatic niche conditions preventing adaptive diversification (Wellborn and Langerhans 2015). By contrast, regions with high species richness might also be assembled by multiple dispersals from nearby regions becoming to be a macroevolutionary sink (Goldberg et al. 2005). In this latter case, the species richness was not build-up by *in situ* speciation mainly but by continued dispersal through time. To evaluate which of these scenarios is more plausible it is necessary to adopt an approach that explicitly infer the number of the dispersal and cladogenetic events across areas (Roy and Goldberg 2007, Dupin et al. 2017).

Differences in species diversification are also considered as a main driver of the geographical diversity gradient for many groups (Kennedy et al. 2014, Pinto-Ledezma et al. 2017). This hypothesis states (Table 2) that differences in net diversification rates between areas are the main driver of differences in regional species richness between areas. Davies and Buckley (2011) used the phylogenetic diversity controlled by species richness (i.e., residual PD -rPD-) to distinguish areas with different evolutionary processes. These authors predicted that areas where rapid speciation and low immigration events from other areas occurred, are dominated by large adaptive radiations (e.g., large islands; Losos and Schluter 2000). By contrast, areas with slow speciation and colonized by multiple lineages through time should have high values of residual PD.

The “out of the tropics” -OTT-hypothesis (Jablonski et al. 2006; Table 2) states that latitudinal diversity gradient is due to that the majority of lineages originated in the tropics and then migrated to extratropical regions. Under this hypothesis, tropics harbour higher net diversification rates (higher speciation and lower extinction) than extratropical regions and dispersal rates are higher from the tropics to extratropical regions than the reverse (Jablonski et al. 2006; Table 1). For instance, Rolland et al. (2014) used the GeoSSE model to test this hypothesis in the generation of the latitudinal mammal diversity gradient. They found that net diversification rates (i.e., the balance of speciation minus extinction) was higher in tropical than in temperate regions and dispersal rates were higher from the tropics to temperate regions than the reverse. Also, Pinto-Ledezma et al. (2017) used the GeoSSE model to test an analogue hypothesis to OTT, as form of Out of the Forest hypothesis (OTF), using Furnariides birds as a clade model. Their favoured a model where open areas have higher speciation, extinction and dispersal rates than forest habitats. All these results suggest that it is reasonable to use either phylogenetic metrics or explicit diversification approaches (e.g., the GeoSSE model) to evaluate a set of evolutionary-based hypotheses as a main driver of geographical diversity gradients. However, we show here (see below) that these approaches fail to capture the evolutionary and biogeographic processes at spatial scales.

### Are phylogenetic metrics capturing well the diversification process across geography?

A deep understanding of evolutionary processes affecting regional species assemblages is coming from the integration of molecular phylogenies and fossil record (Quental and Marshall 2010, Marshall 2017). From this integration of neontological and paleontological perspectives, it is clear that both approaches are necessary to test evolutionary-based hypothesis in macroecological research. Several hypotheses were proposed to explain geographical diversity patterns, particularly the latitudinal diversity gradient -LDG- (see Table 2 for a summary and compilation of the main hypotheses reported in the literature). Although the ideal approach is to generate robust conclusions from multiple lines of evidence (e.g., fossil record, molecular phylogenies, biogeographical inference) it is clear that this information is scarce for many taxonomic groups. Many macroecological studies have adopted either phylogenetic metrics or explicit diversification approaches (e.g., the GeoSSE model) to evaluate the relative contribution of speciation, extinction and dispersal on the resulting geographical diversity gradients (Table S1).

Phylogenetic metrics can be easily visualized in a geographical context and several inferences about ecological (e.g., dispersal) and evolutionary (e.g., speciation) process can be done. As these metrics provide a *per-species level diversification* metric for each species in a phylogeny, it is possible to associate these values with the corresponding species’ geographical range and obtain a mean value for cells or regions in a given geographical domain (Table 1; Figure 2). By contrast, explicit diversification approaches (e.g., GeoSSE; BAMM; fitting models) provide a *per-lineage level diversification* metric for a given clade or a regional assemblage (Rabosky 2016a). However, in some cases, it is possible to generate a *per-species level diversification* metric with these approaches. For instance, Pérez-Escobar et al. (2017) used the function GetTipsRates in BAMMtools (Rabosky et al. 2014) to map speciation rates for Neotropical orchids.

These two approaches (phylogenetic metrics and lineage diversification) potentially can provide complementary pictures about how macroevolutionary dynamics have taken place in the geography. One the one hand, it is possible to estimate diversification rates for a given clade using the number of species, its age and a birth-death models (Magallón and Sanderson 2001, Nee 2006, Sánchez-Reyes et al. 2017). These model-fitting approaches allow to whether diversity- or time-dependent diversification process has taken place in a regional assemblage (Etienne et al. 2012, Rabosky 2014, Valente et al. 2015). On the other hand, per-species diversification rate metrics allow establishing the potential of each individual species to generate more species (Jetz et al. 2012, Rabosky 2014, 2016a). However, these approaches imply at least a different process, which left a different signature on the geography. Phylogenetic metrics captures a total diversification process (Rabosky 2009), whereas lineage diversification approaches (e.g., BAMM) can potentially provide information about an individual diversification process (Rabosky 2013). In addition, still is not clear whether phylogenetic metrics can provide an accurate description of the diversification dynamics across geography.

The first step to clarify how well these phylogenetic metrics behave is to establish a comparison within and between different taxonomic groups. To evaluate how different phylogenetic metrics vary across geography and their relationship with species richness, we used two empirical data sets from our own empirical work (furnariid birds and anole lizards; (Pinto-Ledezma et al. 2017, Velasco et al. 2018) and a data set compiled from several sources (hylid frogs; (Wiens et al. 2006, Algar et al. 2009, Pyron 2014a). We mapped across geography five phylogenetic metrics (Table 1, Figure 2). We selected these three data sets because previous work analysed how evolutionary-based hypotheses affected the present-day species richness gradient (Wiens et al. 2006, Algar et al. 2009, Pinto-Ledezma et al. 2017, Velasco et al. 2018).

Figure 2 shows the geographical pattern of species richness and the five phylogenetic metrics for *Anolis* lizards, hylid frogs and Furnariides birds. For all clades, there is a higher species concentration near to the Ecuador. Higher species concentration for hylids and Furnariides can be found in the Amazon and the Atlantic forest and *for Anolis* lizards in Central America and the Caribe (Figure 2A-C; see also Algar et al. 2009, Pinto-Ledezma et al. 2017, and Velasco et al. 2018, for a detailed description of the geographical species richness pattern for these clades, respectively). In terms of the geographical pattern of each phylogenetic metric (Figure 2D-R), in most of the cases cells with higher metric values are related to cells that contain high species richness and vice versa (Figure 2D-R; Figure A1). However, the degree and the direction of this relationship changes according to the phylogenetic metric used. For example, MRD, a metric of species derivedness, shows a negative correlation with species richness (Figure 2J-L; Figure A1). Importantly, the spatial relationships between species richness and phylogenetic metrics found in our analysis could simply be the result of aggregated species-level attributes within cells or assemblages (Hawkins et al. 2017). Hence, any conclusion derived from these relationships needs to be considered carefully. In addition, there are different levels of correlation between phylogenetic metrics (Figure A1). For example, MDR – MA present a high but negative correlation, and rPD – PSV and MRD – MDR present a mid-high positive correlation (Figure A1). Although there are few studies comparing correlations between metrics (Vellend et al. 2010, Miller et al. 2017), to our knowledge, none previous study compares the similarity of these diversification metrics (Table 1). However, some of these metrics sharing mathematical assumptions, which increase the likelihood of correlation between them. For example, for ultrametric trees, metrics as MDR could be approximated by considering the mean root distance (i.e. MRD metric) from the tips to the root (Freckleton et al. 2008), so further studies exploring the mathematical relation between metrics are needed.

In order to assess if the cells/assemblages on average do not represent a random sampling from the species pool, we applied a simple permutation test to explore the non-randomness in each of the phylogenetic metrics. We applied a null model where the presence-absence matrix (i.e., PAM) was randomly shuffled 1000 times, but maintaining the frequency of species occurrence and observed richness in the cells/assemblages (Gotelli 2000). This kind of null model is standard in studies at the community/assemblage level that use phylogenetic information (Cavender-Bares et al. 2004, 2006). Interestingly, none of the phylogenetic metrics deviates from the null expectation for the three clades (Figure 3). Also, very few cells/assemblages present p-values below the 0.05 threshold, thus indicating that the cells/assemblages present random association among species (Figure 3). These results should be supported by repeating analyses with more clades at different spatial extents, but again, we stress that any result obtained with the use of phylogenetic metrics need to be interpreted carefully.

**Figure 3.**
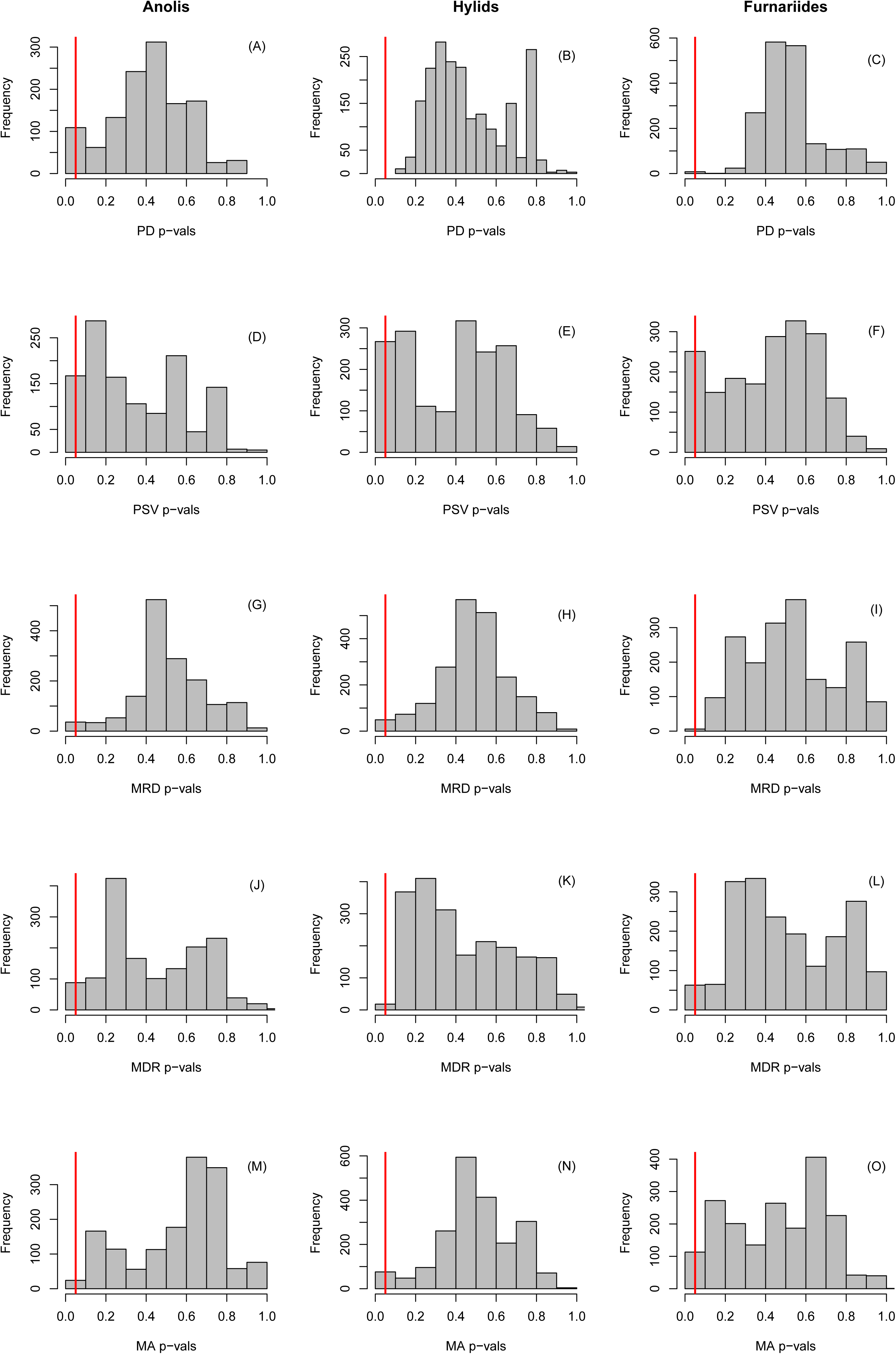
P-values distribution for each phylogenetic metric obtained through the null model (see main text for details). The vertical red lines represent the empirical 0.05 cut-off. Note that for all cases very few cells are below the 0.05 cut-off. (A-C) rPD: residual phylogenetic diversity (i.e., after controlling for species richness); (D-F) PSV: phylogenetic species variability; (G-I) MRD: mean root distance; (J-L) MDR: mean diversification rate; (M-O) Mean ages: average ages of species.

### A brief comparison between phylogenetic metrics and explicit diversification and biogeographic approaches

We compared the phylogenetic metrics enunciated in Table 1, which have been the most used in macroecological research. We explored whether the geographical patterns of these phylogenetic metrics in three empirical examples coincide with the macroevolutionary dynamics inferred using explicit modelling diversification approaches. In particular, we implemented the GeoSSE model to estimate the three parameters (speciation, extinction, and dispersal) between two areas in each taxonomic group (Table 3 and 4). In addition, we used the BAMM approach to generate the per-species level diversification metric implemented in the software BAMM 2.5.0 (Rabosky 2014). In the following section, we discuss each metric and we compare them with the explicit diversification approaches.

**Table 3.** Parameter estimates from the GeoSSE model for three taxonomic groups (Furnariides birds, hylid frogs, and *Anolis* lizards) across two regions. Areas for each taxonomic group as follows: Furnariides birds: A: Forest; B: Open areas; Hylid frogs: A: Extra tropics; B: Tropics; *Anolis* lizards: A: Islands; B: Mainland.

| Group | Rates | A | B | AB |
| --- | --- | --- | --- | --- |
| Furnariides birds | Speciation | 0.139 ± 0.020 | 0.223 ± 0.065 | 0.041 ± 0.020 |
|  | Extinction | 0.040 ± 0.025 | 0.107 ± 0.075 | - |
|  | Dispersal | 0.021 ± 0.004 | 0.311 ± 0.114 | - |
|  | Net diversification | 0.099 ± 0.005 | 0.116 ± 0.01 | - |
| Hylid frogs | Speciation | 0.044 ± 0.003 | 0.044 ± 0.003 | 0.041 ± 0.025 |
|  | Extinction | 0.002 ± 0.002 | 0.002 ± 0.002 | - |
|  | Dispersal | 0.001 ± 0.001 | 0.035 ± 0.010 | - |
|  | Net diversification | 0.042 ± 0.003 | 0.042 ± 0.003 | - |
| *Anolis* lizards | Speciation | 0.058 ± 0.003 | 0.058 ± 0.003 | 1.245 ± 1.303 |
|  | Extinction | 0.001 ± 0.001 | 0.001 ± 0.001 | - |
|  | Dispersal | 0.002 ± 0.001 | 0.0003 ± 0.000 | - |
|  | Net diversification | 0.057 ± 0.002 | 0.057 ± 0.002 | - |

**Table 4.** Frequency of dispersal events inferred using biogeographical stochastic mapping (BSM) for three taxonomic groups (Furnariides birds, hylid frogs, and *Anolis* lizards) across two regions. Areas for each taxonomic group as follows: Furnariides birds: A: Forest; B: Open areas; Hylid frogs: A: Extra tropics; B: Tropics; *Anolis* lizards: A: Islands; B: Mainland.

| Event | Group | Regions | A | B |
| --- | --- | --- | --- | --- |
| Range expansions | Furnariides birds | A | 0 | 92.62 ± 4.39 |
|  |  | B | 31.4 ± 4.29 | 0 |
|  | Hylid frogs | A | 0 | 0.64 ± 0.78 |
|  |  | B | 12.92 ± 0.88 | 0 |
|  | *Anolis* lizards | A | 0 | 0 |
|  |  | B | 0 | 0 |
| Founder events | Furnariides birds | A | 0 | 24.46 ± 2.54 |
|  |  | B | 10.88 ± 2.22 | 0 |
|  | Hylid frogs | A | 0 | 0.66 ± 0.66 |
|  |  | B | 4.3 ± 1.42 | 0 |
|  | *Anolis* lizards | A | 0 | 2.02 ± 0.14 |
|  |  | B | 0.12 ± 0.33 | 0 |

#### Residual Phylogenetic Diversity (rPD)

In the case of furnariid birds, we show that forest areas tend to exhibit slightly higher values of rPD in contrast with open areas (Figure 4; see also Figure 2). According to Davies and Buckley’s logic, these areas exhibit slow diversification and frequent dispersal from open areas. Pinto-Ledezma et al. (2017) using GeoSSE and BAMM approaches indicated that open areas exhibit higher net diversification rates than open areas (Table 3). For hylid frogs, we found that tropical areas tend to exhibit higher rPD values than extratropical regions (Figure 5). However, by adopting an explicit diversification approach (GeoSSE and BAMM), we found that net diversification rates were similar in both regions (Table 3). In the case of *Anolis* lizards, the rPD values were higher in the continent than in the island areas (Figure 6). However, using GeoSSE and BAMM, we found that both rates were similar (Table 3). In a recent paper, Poe et al. (in press) also found that macroevolutionary rates are similar between insular and mainland clades. All these results suggest that rPD likely does not provide an accurate signature of the macroevolutionary dynamic at spatial scales. In fact, it seems that rPD tends to overestimate differences between regions when a stationary diversification process is occurring across geography. A potential solution might be rethinking the way in which we visualize rPD across geography in contrast with the original meaning by Davies & Buckley (2011; see also Forest et al. 2007).

**Figure 4.**
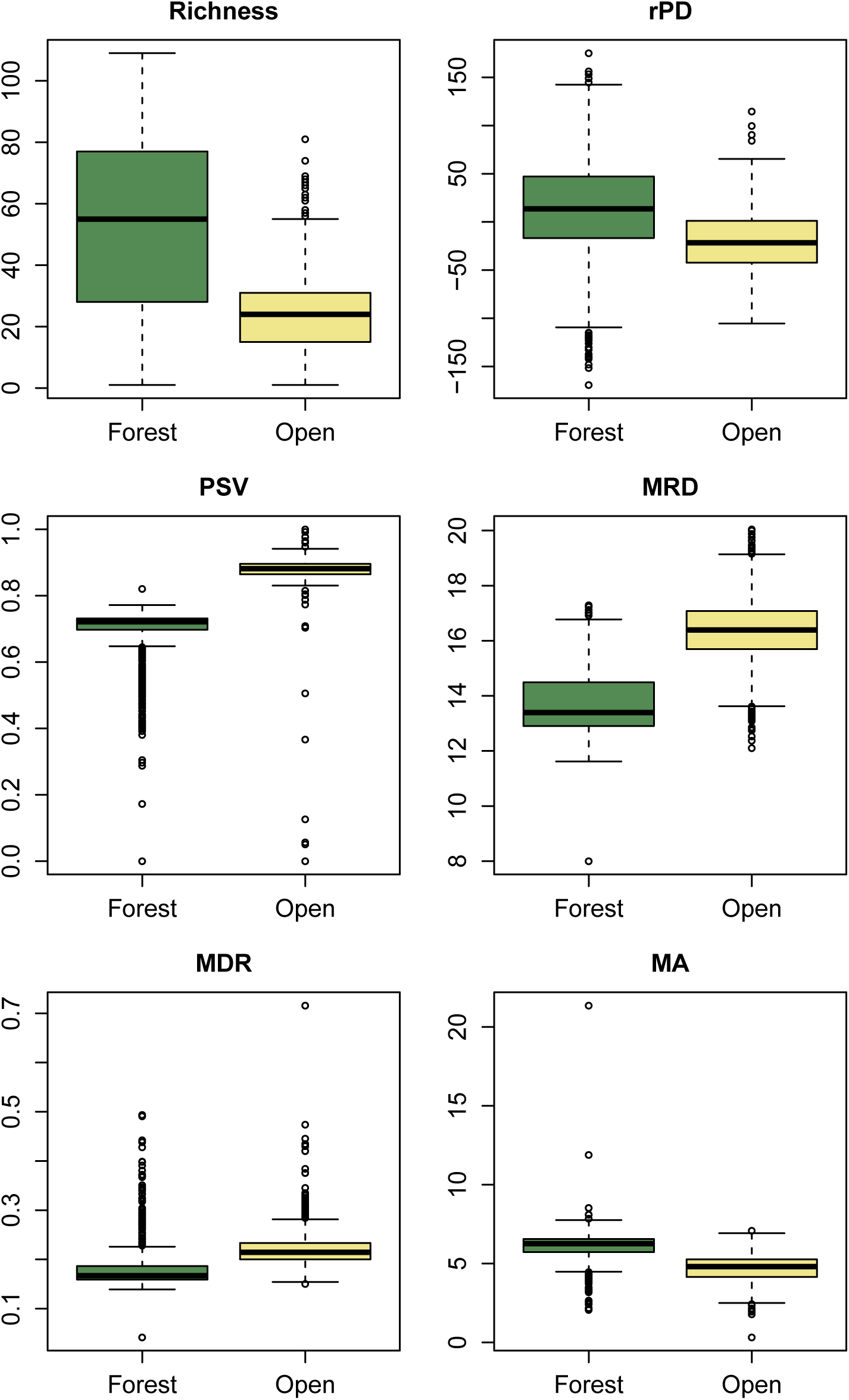
Variation of phylogenetic metric values for Furnariides birds in forest and open areas. rPD: residual phylogenetic diversity (i.e., after controlling for species richness); PSV: phylogenetic species variability; MRD: mean root distance; MDR: mean diversification rate; Mean ages: average ages of species.

**Figure 5.**
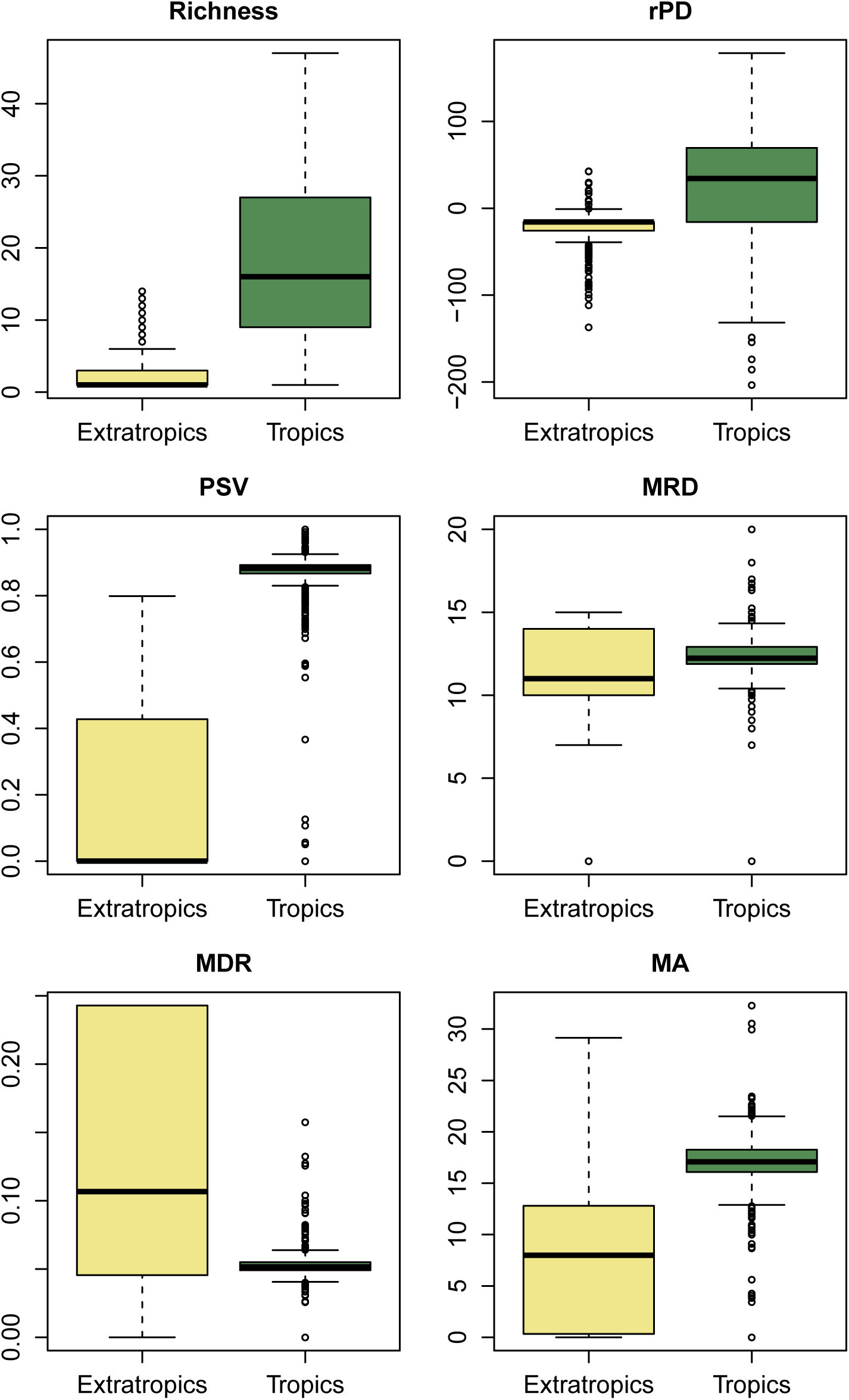
Variation of phylogenetic metric values for Hylid frogs in tropics and extra-tropics regions. rPD: residual phylogenetic diversity (i.e., after controlling for species richness); PSV: phylogenetic species variability; MRD: mean root distance; MDR: mean diversification rate; Mean ages: average ages of species.

**Figure 6.**
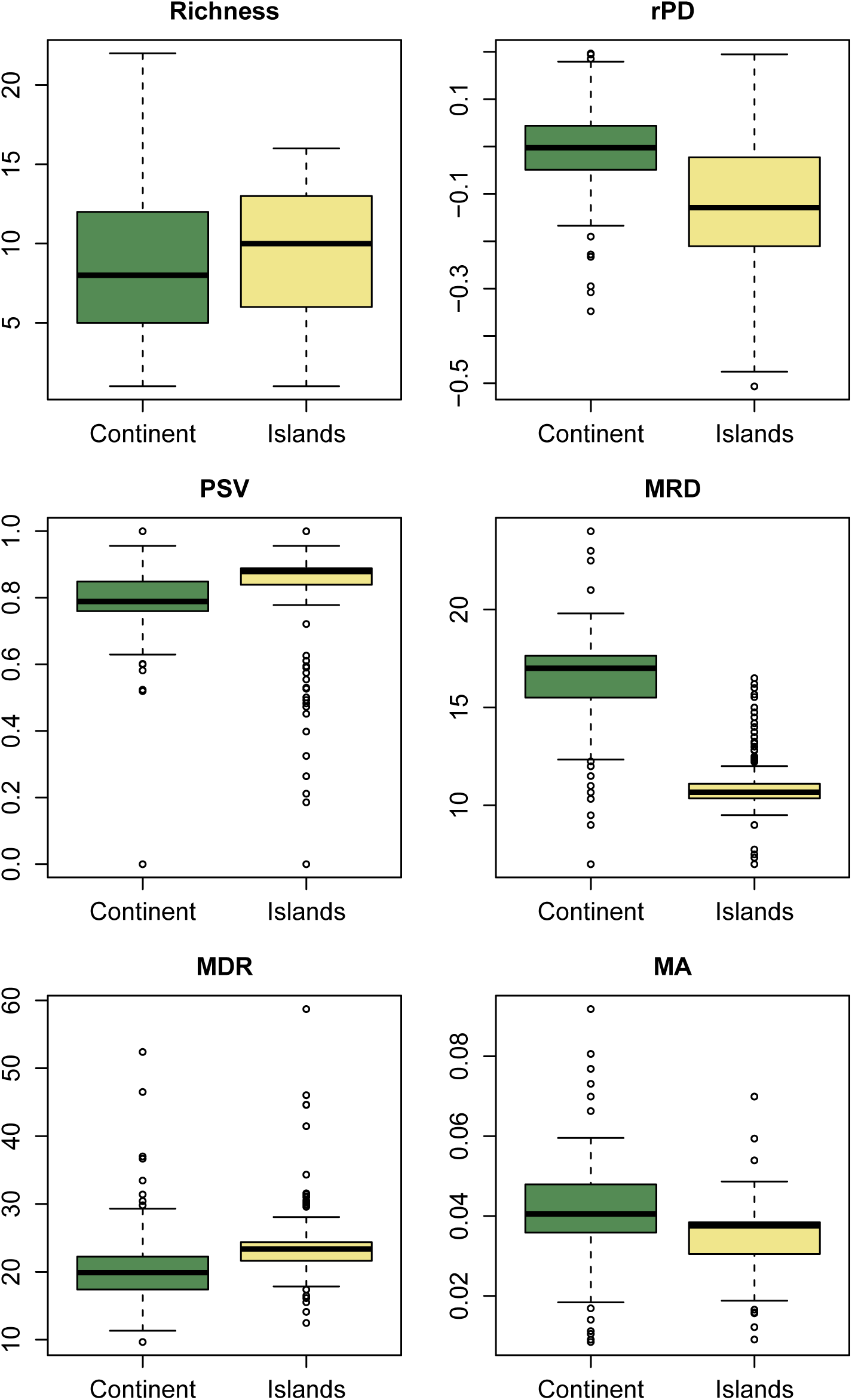
Variation of phylogenetic metric values *for Anolis* lizards in continental and insular areas. rPD: residual phylogenetic diversity (i.e., after controlling for species richness); PSV: phylogenetic species variability; MRD: mean root distance; MDR: mean diversification rate; Mean ages: average ages of species.

#### Mean root distance (MRD)

As we discussed, *MRD* captures a total diversification value portraying the number of cladogenetic events co-occurring in a given region. In the case of furnariid birds, we found that MRD values tend to be higher in open than forest areas (Figure 4). Accordingly, this metric suggests that more cladogenetic events were accumulated in open areas (i.e., more total diversification; Rabosky 2009). Therefore, this metric, for this bird clade, is consistent with results from explicit diversification approaches (Table 3). For hylid frogs, it seems that there are no differences in MRD values between extratropics and tropics areas (Figure 5). However, tropical areas have some cells with very high values. Again, MRD provide an accurate description of the total diversification pattern in this clade across the latitudinal gradient. In *Anolis* lizards, we found that MDR values tend to be lower in islands in comparison with mainland areas (Figure 6). In this case, MRD did not provide an accurate description of the evolutionary processes occurring between the mainland and insular anole assemblages. However, there is also a high probability that the high MRD values in the mainland are a direct reflect of an idiosyncratic evolutionary trajectory of each one of the two clades that radiated there (i.e., *Draconura* and *Dactyloa* clade; see Poe et al. 2017, Velasco et al. 2018). These two clades seems to exhibit differential diversification dynamics across geography (Velasco et al. 2018) but further research might be necessary to evaluate these differences.

#### Phylogenetic species variability (PSV)

The PSV metric provides information about how related are the species in a given regional assemblage. In hylid frogs, we found that tropical assemblages tend to be composed of more related species than extratropical assemblages (Figure 5). Hylid assemblages in extratropical areas are composed of multiple lineages that dispersed from tropical areas and then diversified there. We found higher dispersal rates from tropical to temperate regions than vice versa (Table 3 and 4). The same tendency is present in the case of furnariid birds where open areas exhibit higher PSV values than forest areas (Figure 4) and dispersal rates were higher from open to forest areas than the reverse (Table 3). By contrast, we did not find any evidence for differences in PSV values between island and mainland *Anolis* assemblages (Figure 6). In addition, the dispersal rates were very low between these two regions (Table 3; Poe et al. 2017). All these results confirm that the PSV metric can provide some insights about how dispersal process have shaped regional assemblages. We find evidence that low PSV values (i.e., phylogenetically over-dispersed faunas) are influenced by multiple dispersals along its evolutionary history.

#### Mean diversification rate (MDR)

Jetz et al. (2012) proposed MDR metric as a species-level speciation rate metric based in the branch length along the path from the root of a tree to each individual species. In furnariid birds, we noted that MDR was slightly higher in open versus forest areas and the same pattern is present using the BAMM approach (Pinto-Ledezma et al. 2017; Figure 5; Table 3). For hylid frogs, extratropical regions tend to exhibit higher values than tropical regions (Figure 5). MDR seems to capture well the differences in macroevolutionary diversification for these taxa along the latitudinal diversity gradient. A similar pattern is present when the BAMM approach is used (Table 3). We consider that both metrics (MDR vs per-species diversification rate from BAMM) leave the same signature in the geography. In *Anolis*, we found that insular assemblages tend to exhibit higher MDR values than continental assemblages (Figure 6), However, there is no difference in the macroevolutionary dynamic between these two areas for the *Anolis* lizards clade (Poe et al. in press, Velasco et al. 2018).

#### Mean ages (MA)

The average of ages of co-occurring lineages are used to test evolutionary hypothesis about whether a region maintains older lineages than others (e.g., a museum) or a combination of old and recent lineages (e.g., OTT hypothesis, Table 1). Although this metric does not provide any inference of the ancestral area of the clade, it is possible to implement an explicit biogeographic approach to test this (see below). For example, in hylid frogs, we found that extratropical areas are composed of older lineages than tropical regions (Figure 5). The biogeographic parametric approach infers this same area as ancestral for the entire lineage (Figure A2). In furnariid birds, mean ages metric revealed that older lineages have accumulated more in forest than open areas (Figure 4). In accordance, the ancestral area inferred with a parametric biogeographic method was the forest area (Figure A3). In the case of the anole lizards, insular settings tend to be composed of older lineages than continents. However, the ancestral area for the entire anole clade is the mainland, particularly South America (Poe et al. 2017). The mainland *Anolis* radiation is composed of two clades, one clade that originated in South America (the *Dactyloa* clade; Poe et al. 2017) and colonized Caribbean islands, and the other clade (the *Norops* clade; Poe et al. 2017) that originated in the Caribbean islands and then colonized back the mainland in Middle America and then dispersed to South America. Therefore, the biogeographical history of *the Anolis* radiation is complex and involves multiple dispersals between islands and mainland areas (Poe et al. 2017; Figure A4). In general, mean ages does not provide enough information about the biogeographic origin and maintenance of a clade. This happens because multiple dispersals and *in situ* cladogenesis might erase any simplistic pattern elucidated for this metric, as found in the case of *the Anolis* lizards.

## How Dispersal and Extinction Affect Inferences of Geographical Diversification Gradients?

Dispersal is another key macroevolutionary process that ultimately determines the number of a species in a region (Roy and Goldberg 2007, Eiserhardt et al. 2013, Rolland et al. 2014, Chazot et al. 2016). However, few studies evaluated explicitly how the direction of dispersals between region contributes to the generation of regional differences between areas (Chown and Gaston 2000, Goldberg et al. 2005, 2011, Jablonski et al. 2006). Roy and Goldberg (2007) showed with simulations that dispersal asymmetry between areas had a strong impact in the regional species richness and the average age of these lineages. Accordingly, phylogenetic metrics can be sensitive to dispersal between areas because it is impossible to distinguish which lineages originated by *in situ* speciation or simply due to dispersal from nearby areas. Goldberg et al. (2011) developed the GeoSSE model to evaluate how range evolution affected diversification rates in a phylogenetic comparative approach. The GeoSSE model only considers three states (A: endemic species to a region; B: endemic species to another region; and AB for widespread species) and makes a series of assumptions that can be problematic. The first assumption of the GeoSSE model is that a time-dependent process dominates the diversification dynamic in each region (Stephens and Wiens 2003, Wiens 2011). This assumption conflicts with a diversity-dependent process assumption and this debate is far from being resolved (Cornell 2013, Harmon and Harrison 2015, Rabosky and Hurlbert 2015). The second problematic assumption has to do with the fact that the GeoSSE model consider dispersal rates as stable through time and lineages. In other words, the dispersal ability and therefore the frequency of transitions between areas are constant across the evolutionary history of a clade. There are many empirical evidence showing that dispersal rates vary across time and space among lineages (McPeek and Holt 1992, Sanmartín et al. 2008, Robledo-Arnuncio et al. 2014).

Regardless of these major assumptions, the GeoSSE model has been adopted to evaluate relative contributions of speciation, extinction and dispersal to the generation of species richness gradients (e.g., Rolland et al. 2014, Pyron 2014b, Staggemeier et al. 2015, Looney et al. 2016, Morinière et al. 2016, Pulido-Santacruz and Weir 2016, Alves et al. 2017, Hutter et al. 2017, Pinto-Ledezma et al. 2017). In a recent study, Rabosky and Goldberg (2015) found that state-dependent diversification models tend to inflate excessively the false discovery rates (i.e., type I error rates). In particular, Rabosky and Goldberg (2015) found that these models tend to find false associations between trait shifts and shifts in macroevolutionary dynamics. Although the Rabosky and Goldberg’s study was not based on the GeoSSE model, it is clear that transitions between areas (i.e., dispersal events) can be falsely associated with shifts in speciation and extinction rates across the phylogeny. Alves et al. (2017) also found that geographical uncertainties in the assignment of species to a given area affect the parameter estimates (i.e., speciation, extinction and dispersal rates) in the GeoSSE model. Same authors also evaluated how incorrect assignments of bat species to tropical or extra-tropical regions can generate erroneous conclusions about the relative role of speciation, extinction and dispersal on a latitudinal diversity gradient. From these studies, it is clear that dispersal is a major issue that needs to be evaluated explicitly in macroecological studies.

Pulido-Santacruz and Weir (2016) also used the GeoSSE model to disentangle the relative effect of speciation, extinction and dispersal on the latitudinal bird diversity gradient. They found that extinction was prevalent across all bird clades and therefore they suggest this as a main driver of the geographical bird diversity gradient. Pyron (2014c), also using the GeoSSE model, found that temperate diversity in reptiles is due to higher extinction in these areas. We consider that extinction inferences from the GeoSSE model should be treated with caution. For the few clades where fossil record is abundant (e.g., marine bivalves; Jablonski et al. 2006), studies point out to conclude that extinction differences between regions should be treated with caution due to the potential sampling bias (Jablonski et al. 2006, 2017). In addition, studies based on extensive simulations found that extinction inferences based only in molecular phylogenies are not reliable (Rabosky 2010a, 2016b, Quental and Marshall 2010), although extinction rates can be estimated relatively well using medium to large phylogenies (Beaulieu & O’Meara 2015).

In a recent review, Sanmartín and Meseguer (2016) proposed that it is possible to detect the extinction signature in molecular phylogenies using extensive simulations and lineage-through-time -LTT-plots (see also Antonelli and Sanmartín 2011). These authors also found that many birth-death models leave a similar phylogenetic imprint, which make indistinguishable some scenarios. In addition, extinction events can affect substantially the ancestral range estimates, and therefore dispersal and extinction parameters in several parametric biogeographic methods (e.g., Dispersal-Vicariance –DIVA- and Dispersal-Extinction-Cladogenesis –DEC-models; Ronquist 1997, Ree et al. 2005). Sanmartín and Meseguer (2016) finally proposed that the adoption of a hierarchical Bayesian approach using continuous-time Markov Chain models will allow a better estimation of extinction both in geography and in the phylogeny (Sanmartín et al. 2008, Sanmartin et al. 2010).

Recently, Rabosky and Goldberg (2017) developed a semi-parametric method (FiSSE) to correct the statistical problems found in BiSSE models by themselves in a previous paper (Rabosky and Goldberg 2015). However, the FiSSE method does not allow the evaluation of the contribution of dispersal on regional species richness. In any case, the best suitable framework to estimate relative contributions of speciation, extinction and dispersal might be the GeoSSE model (or parametric biogeographic models; e.g., Matzke 2014; see below), although it requires the simulation of a series of null scenarios to evaluate the statistical power in each case (see Alves et al. 2017, Pinto-Ledezma et al. 2017 for a few examples). For instance, Pinto-Ledezma et al. (2017) developed a parametric bootstrapping approach simulating traits to evaluate whether empirical inferences are different from the simulated. They simulated 100 datasets of neutral characters along a set of empirical phylogenies and using this new information repeated the same procedure with empirical data (see Appendix S1 in Pinto-Ledezma et al. 2017 for details of the bootstrapping approach). This bootstrapping procedure assumes no direct effect of the geographic character states on the parameter estimations (Feldman et al. 2016, Pinto-Ledezma et al. 2017).

Finally, it should be clear that more research would be necessary to establish how extinction affect estimation parameters in state-dependent diversification approaches (e.g., the GeoSSE model). For instance, the inclusion-exclusion of extinct species in simulated phylogenies using birth-death models could substantially affect the geographical inferences of speciation, extinction and dispersal parameters in the GeoSEE model. This kind of approach might provide some lights on how to biased can be the parameter estimates with only molecular phylogenies using the GeoSSE model or any other modeling approach.

### Parametric biogeographical approaches in macroecological studies

The use of parametric biogeographic approaches is an optimal solution to estimate dispersals across time and space (Matzke 2014, Dupin et al. 2017). These methods are promising in identifying the relative roles of cladogenetic and anagenetic processes shaping regional species richness. Recently, Dupin et al. (2017) developed a biogeographical stochastic mapping to infer the number of dispersals, and other biogeographical events, in the evolutionary history of Solanaceae plants across the world. This approach allows the inference from multiple process including sympatric speciation, allopatric speciation, founder-event speciation, range expansion (i.e., dispersal without speciation) and local extinction (i.e., range contractions) based on a time-calibrated phylogenetic tree and the occurrence of species in geographical regions (see also Matzke 2014 for more detailed description of the method). These explicit biogeographical approaches are promising in macroecological studies since they allow to test simultaneously a set of evolutionary process during the diversification of a clade in a region (Velasco 2018). In addition, with these new approaches it is possible to differentiate effectively between macroevolutionary sources and sink areas (Goldberg et al. 2005, Castroviejo-Fisher et al. 2014, Poe et al. 2017). For instance, Poe et al. (2017) used a parametric biogeographical approach to estimate the number of events among regions and distinguish those areas where many cladogenetic events occurred (i.e., *in situ* speciation) and areas where almost all its diversity was build-up from extensive colonization of other regions.

The biogeographical stochastic mapping (BSM) method developed by Dupin et al. (2017) is promising to estimate more accurately the number of dispersal events between regions based on a better estimation of the ancestral area for a clade. We evaluated how dispersal rates between regions can affect inferences drawn only from phylogenetic metrics in our three data sets. We implemented GeoSSE and BSM approaches for each data set (Table 3 and 4). For the case of hylid frogs, we counted the inferred number of dispersal events between tropical and extra-tropical regions in the Americas (Table 3; see also (Wiens et al. 2006, Algar et al. 2009). For furnariid birds, we counted the number of dispersal events between open and forest areas (Table 3; see also Pinto-Ledezma et al. 2017). Finally, for anole lizards, we counted the number of dispersal events between insular and mainland areas (Table 3; see also Algar and Losos 2011, Poe et al. 2017, Velasco et al. 2018).

Using biogeographical stochastic mapping -BSM-, we inferred the number of dispersal events from one region to another for each one of the three taxonomic groups examined (Table 3). The BSM approach allows us to disentangle which dispersals were only range expansions and which dispersals generated a speciation event (i.e., a founder-event speciation; (Barton and Charlesworth 1984, Templeton 2008). For furnariid birds, we found that range expansions were three times higher from forest areas to open areas than the reverse and founder events were twice higher from forest to open areas than the opposite (Table 3). This result suggests that differences in species richness between forest and open areas are due by recurrent dispersal events along the furnariid diversification history (Pinto-Ledezma et al. 2017). Pinto-Ledezma et al. (2017) found a similar result using the GeoSSE approach, but they conducted a parametric simulation approach to evaluate whether there was a direct effect of the geographic location on the parameter estimates. Their results show that the GeoSSE approach, in this case, had limited power to detect a signature of geographic region on speciation, extinction and dispersal rates. With the implementation of the BSM approach here, we corroborate Pinto-Ledezma et al.’s findings with improved statistical power. In the case of hylid frogs and the transitions between tropical and extra-tropical areas, we found that the BSM approach inferred more dispersal events from tropical to extra-tropical regions (Table 3 and 4). However, the number of founder events was relatively low in comparison with range expansions (Table 3). These results suggest that few dispersal events have occurred across the diversification of hylid frogs and corroborate that the species richness in each region largely originated by *in situ* speciation modulated by climatic factors (Wiens et al. 2006, Algar et al. 2009). Finally, *for Anolis* lizards, we found that dispersal events between insular and mainland regions were relatively low (Table 3 and 4). We did not find evidence of any expansion range events from mainland to island or vice versa. This also corroborates previous findings that evolutionary radiation of anole in insular and mainland settings is due to extensive *in situ* diversification (Poe et al. in press, 2017, Algar and Losos 2011).

These results point out that the BSM approach (Dupin et al. 2017) is a promising approach when we are interested in testing the role of anagenetic and cladogenetic events on the resulting geographical species richness gradients. Although parametric biogeographic approaches are still in their infancy (Sanmartín 2012, Matzke 2014, Dupin et al. 2017), these methods allow us to evaluate macroevolutionary dynamics (i.e., speciation and extinction) in an explicit geographical context. These methods are statistical powerful and make use of a series of explicit geographic range evolution models (Velasco 2018).

## Toward an Integration of Biogeographical and Species Diversification Approaches in Macroecological Studies

Although different parametric biogeographic methods have been developing at least for the last 20 years (Ronquist 1997, Ree et al. 2005, Landis et al. 2013, Matzke 2014, Dupin et al. 2017), the adoption of these methods to test evolutionary-based hypotheses underlying geographical diversity gradient has been rare. For instance, few studies examined here tested the effect of dispersal events in the generation of regional species richness assemblages. It should clear that the current paradigm in biogeography makes a call for an evaluation of the relative frequency of cladogenetic and anagenetic process during the biogeographical history of lineages. The adoption of parametric approaches in future macroecological studies will contribute to an improvement of the estimation of speciation, extinction and dispersal processes as drivers of the geographical diversity gradients. In addition, we also think that it is necessary to establish which macroevolutionary dynamics govern regional assemblages. Phylogenetic approaches based on fitting diversification models help to test whether regional species richness is due to diversity dependence (i.e., ecological limits), time dependence, or environmental factors (Rabosky and Lovette 2008, Etienne et al. 2012, Etienne and Haegeman 2012, Condamine et al. 2013). We also stress that the adoption of many approaches providing multiple lines of evidence will help to disentangle the evolutionary and ecological causes of biodiversity gradients. Some recent studies have pointed toward this strategy and have begun to provide evidence from many lines to understand how evolutionary processes underlying species richness gradients works (Hutter et al. 2017, Pinto-Ledezma et al. 2017).

### Conclusions and recommendations

The resulting geographical pattern of several phylogenetic metrics did not provide any robust evidence of a spatially explicit diversification dynamic. As we have shown, these resulting geographical patterns did not differ from that generated by a simple null model. It is hard to untangle causal mechanisms (i.e., speciation, extinction, and dispersal) from only the geographical signature that these metrics attempt to capture. We recommend that phylogenetic metrics should be used only to visualize geographical patterns of total diversification (e.g., MRD, residual PD; MDR), phylogenetic structure (e.g., PSV), or mean ages of co-distributed species (e.g., MA) (Table 1). We suggest that conclusions about the role of evolutionary processes in the generation and maintenance of species richness gradients based only in these phylogenetic metrics should be avoided and additional approaches always should be used.

Some explicit diversification approaches (e.g., model fitting approaches; (Etienne et al. 2012, Rabosky 2014, Valente et al. 2015) are useful to establish the macroevolutionary dynamics operating at regional scales. Although some approaches (e.g, the GeoSSE model) allow us to evaluate the relative role of the ultimate process that modify the regional species diversity, its statistical power (e.g., high Type I errors) has been challenged by simulation and empirical studies. Furthermore, the extinction and dispersal estimates inferred by the GeoSSE model tend to be unbiased. Parametric biogeographic approaches are becoming a standard tool to evaluate how evolutionary processes can explain the geographical distribution of extant taxa. These approaches are promising and should be extensively used because allow us to estimate the relative frequency of cladogenetic and anagenetic process shaping the regional species richness. It is necessary that macroecological studies use a combination of explicit diversification approaches and parameter biogeographic methods with the aim to clarify how evolutionary process have shaped regional species richness assemblages. As Jablonski et al. (2017) have outlined, one of the main obstacles to generate an appropriate understanding of the causal mechanisms underlying geographical diversity gradients has been that many studies have tested a single hypothesis, either evolutionary or ecological, as an explanatory factor. We suggest that ecological and evolutionary hypotheses should be tested simultaneously to explain the relative contribution of each process to the regional diversity. As shown by our empirical comparison of phylogenetic metrics, explicit diversification models, and historical biogeographic methods have showed, it is necessary to obtain evidence of different approaches to guarantee sound conclusions about the evolutionary causes of these biodiversity gradients.

## Acknowledgments

We thank Josep Padullés for reviewing the English. JAV is supported by a Postdoctoral fellowship from DGAPA program at Facultad de Ciencias of the UNAM. JAV thanks to Oscar Flores Villela for providing a working space at Museo of Zoologia “Alfonso L. Herrera” and insightful discussions about historical biogeography and evolutionary biology. JNPL is supported by a Postdoctoral research fellowship from the Grand Challenge in Biology Postdoctoral Program, College of Biological Sciences, University of Minnesota.

